# Response to “No support for the adaptive hypothesis of lagging-strand encoding in bacterial genomes”

**DOI:** 10.1101/2020.05.26.117366

**Authors:** Houra Merrikh, Christopher Merrikh

## Abstract

Several previous studies by our group suggest that positive selection can drive certain (not all) genes to be retained in the lagging strand orientation^2,5,6^. This is likely the result of multiple factors including accelerated evolution through replication-transcription conflicts. Liu and Zhang challenge this view, and claim that our method for detecting gene inversions is flawed. Below, we provide empirical evidence that their criticisms are largely unfounded, and show that our original analysis and conclusions are well supported. Though the GC skew method does have a detection limit, we provide new evidence that the fundamental assumptions of our model are accurate, and introduce an improved GC skew calculation which correctly identified 100% of the authors’ gene inversions. Our new findings indicate that the trends we originally identified are stronger than they initially appeared: across species, 89-96% of lagging strand genes appear to be natively leading strand genes that changed orientation. Our statistical analyses offer further support for the notion that for some genes, the lagging strand orientation can be adaptive.

---

Inconsistent vocabulary is a fundamental source of confusion in this debate. Liu and Zhang’s phylogeny-based method identifies conserved orthologs that changed orientation during the divergence an ancestor and descendant species. The authors refer to all such events as “inversions” in accordance with common usage. However, this is distinct from the definition we used^1^. To clarify the language, we will refer to all change-of-orientation events as “flips”. There are two flip sub-types: The first results cause a gene to have negative GC skew (Figure 1, upper graphs). We originally called these “inversions”. However, for clarity we now call them “GC skew inversions”. In the second sub-type, the flip results in a positive GC skew (Figure 1, lower graphs). Here we introduce the term “GC skew reversions” as these events appear to place a gene back in its typical evolutionary orientation. Our method cannot, and was never intended to identify GC skew reversions. Thus, our manuscript’s “inversions” (GC skew inversions) should represent only a subset of Liu and Zhang’s “inversions” (flips).

**Figure 1.**
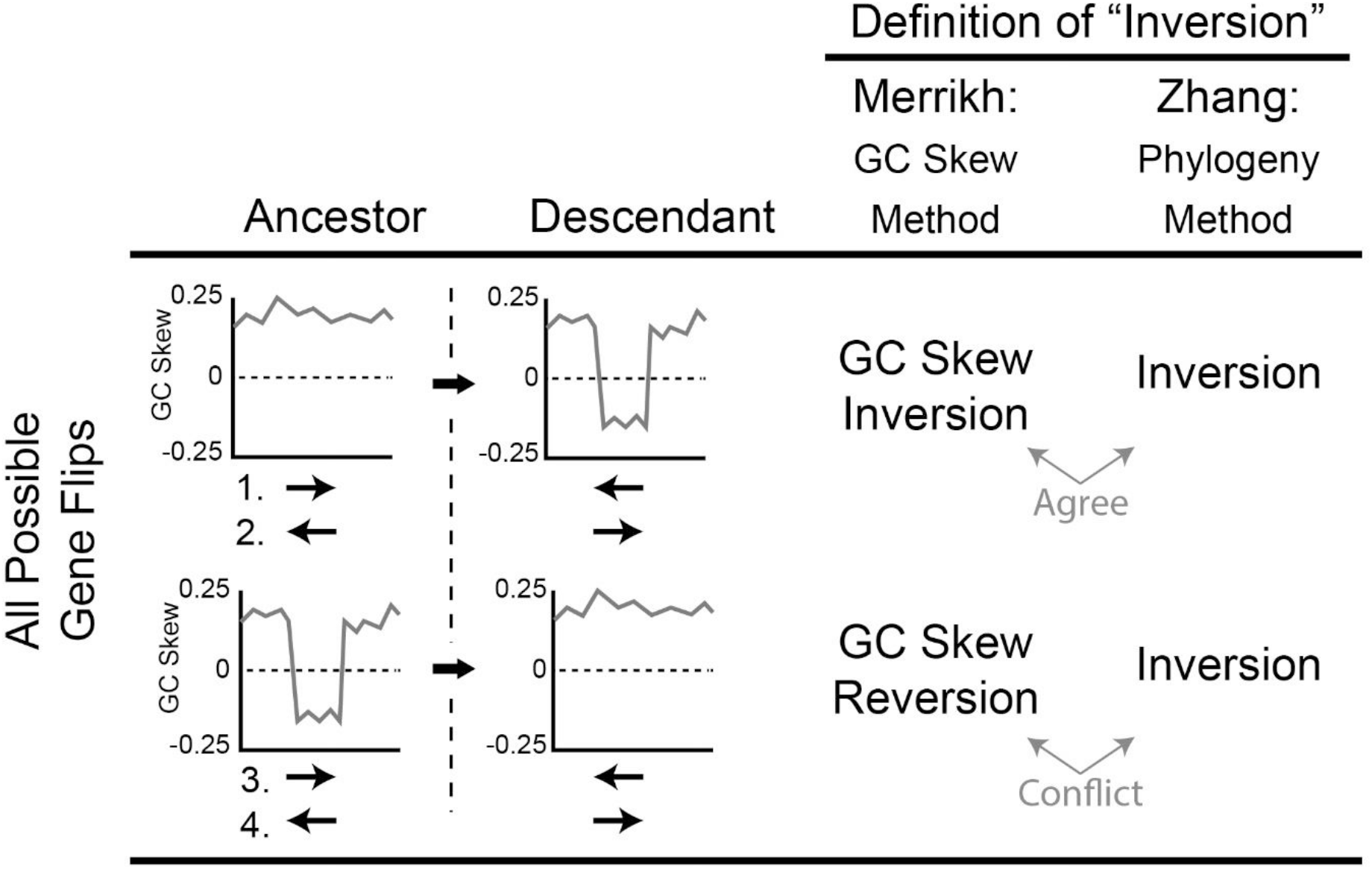
Comparison of gene “inversion” definitions and detection methods. The two studies use different definitions of “gene inversion”. For clarity, we use “Gene Flip” to mean any change-of-orientation event. Gene flips have two subtypes resulting in a gene having a negative GC Skew (upper graphs) or a positive GC skew (lower graphs). The Merrikh and Zhang definitions agree in the first scenario (upper graphs), but conflict in the second (lower graphs). As both gene orientation and GC skew sign (+/−) are important considerations, there are four possible circumstances (numbered 1-4) indicated under the GC skew plots. In case 1, a natively leading strand gene (indicated by the positive GC skew) undergoes a flip resulting in a GC skew inversion. In 2, a natively lagging strand gene undergoes a flip, also resulting in a GC skew inversion. In 3, a leading strand gene with a negative GC skew flips, resulting in a GC skew reversion. In 4, a lagging strand gene with a negative GC skew flips, resulting a GC skew reversion.

To investigate the source of the inconsistencies between the two studies, we calculated the GC skew for each ancestor/descendant orthologue Liu and Zhang identified (this manuscript’s Supplementary Tables 1, 2, and 3). These data show that many flips are GC skew inversions, while others are GC skew reversions. This confirms that the differences between the authors’ data and our own are entirely appropriate, and that most of the “false negative errors” are simply GC skew reversion events (lines 105-106). For example, in the *M. penetrans*/*M. gallisepticum* comparison, the GC skew indicates that 27/52 flips are GC skew inversion and 22/52 are GC skew reversion events, thus, explaining 88% (22/25 total) of the perceived errors. Interestingly, in line 104, the authors report a particularly low agreement between the two methods for lagging-to-leading strand flips. This is because most lagging-to-leading strand gene flips are GC skew reversions (Supplementary Tables 1, 2, and 3).

Liu and Zhang also suggest improving our method’s accuracy by using only 3^rd^ codon position nucleotides, rather than the whole gene sequence when calculating the GC skew (lines 77-78). We hypothesized that third codon position (CP3) nucleotides should be the least reliable source of information because they can mutate without significant consequence. Therefore, after a GC skew inversion, the CP3-based GC skew value should rapidly rise, preventing detection. An additional consideration is that the positive GC skew of whole chromosome arms is clearly detected using a sliding window which completely ignores codon position^2^. As such, modification seemed unnecessary. Nevertheless, we tested the authors’ hypothesis by calculating the GC skew using either the whole gene sequence, or only nucleotides in the 1^st^, 2^nd^, or 3^rd^ codon positions (Table 1). We observe good agreement between the whole gene and the codon position and 2 analyses. However, as predicted, there is lower agreement between the whole gene and 3^rd^ codon position-based calculations, especially for lagging strand genes (Table 1, and our original Figure 5). As we previously showed, the lagging strand is enriched in GC skew inversions, explaining this observation^1^. Hence the 3^rd^ codon position-based GC skew calculation is the lowest fidelity method for GC skew inversion detection.

**Table 1.** Comparison of GC Skew calculation methods. Pearson correlation coefficient values are shown for GC skew values calculated using the whole gene average versus only nucleotides in codon positions 1, 2, or 3. Correlation coefficients are for all leading or lagging strand genes in the indicated species. The lowest correlation is typically between the whole gene average and codon position 3 method for head-on genes. This observation directly supports the validity of the whole-gene average GC skew calculation method used in our original manuscript.

| GC Skew:<br>Calculated for<br>Full Gene<br>versus: | Pearson Correlation Coefficients |  |  |  |  |  |
| --- | --- | --- | --- | --- | --- | --- |
|  | *M. gallisepticum* |  | *B. subtilis* |  | *S. aureus* |  |
|  | Lead. | Lagg. | Lead. | Lagg. | Lead. | Lagg. |
| Codon Pos. 1 | 0.54 | 0.32 | 0.51 | 0.55 | 0.58 | 0.65 |
| Codon Pos. 2 | 0.51 | 0.59 | 0.43 | 0.47 | 0.54 | 0.58 |
| Codon Pos. 3 | 0.44 | 0.07 | 0.53 | 0.45 | 0.42 | 0.35 |

Interestingly, both CP1 and CP3-based GC skew values are generally positive, matching the overall trend (Figure 2A, Table 1). In fact, the magnitude of the GC skew is greatest for CP1 nucleotides (Fig 2. Top graphs). As CP1 nucleotides are under maximal selection pressure (relative to CP2 or CP3), these data suggest that the GC skew values appear to be more strongly influenced by replication related mutation bias than selection (line 74). Additionally, as negative values are almost exclusively associated with lagging strand genes, gene flipping (GC skew inversions) appears to be the best explanation for negative CP1-based GC skew values. Both observations support the validity of our interpretations/methods.

**Figure 2.**
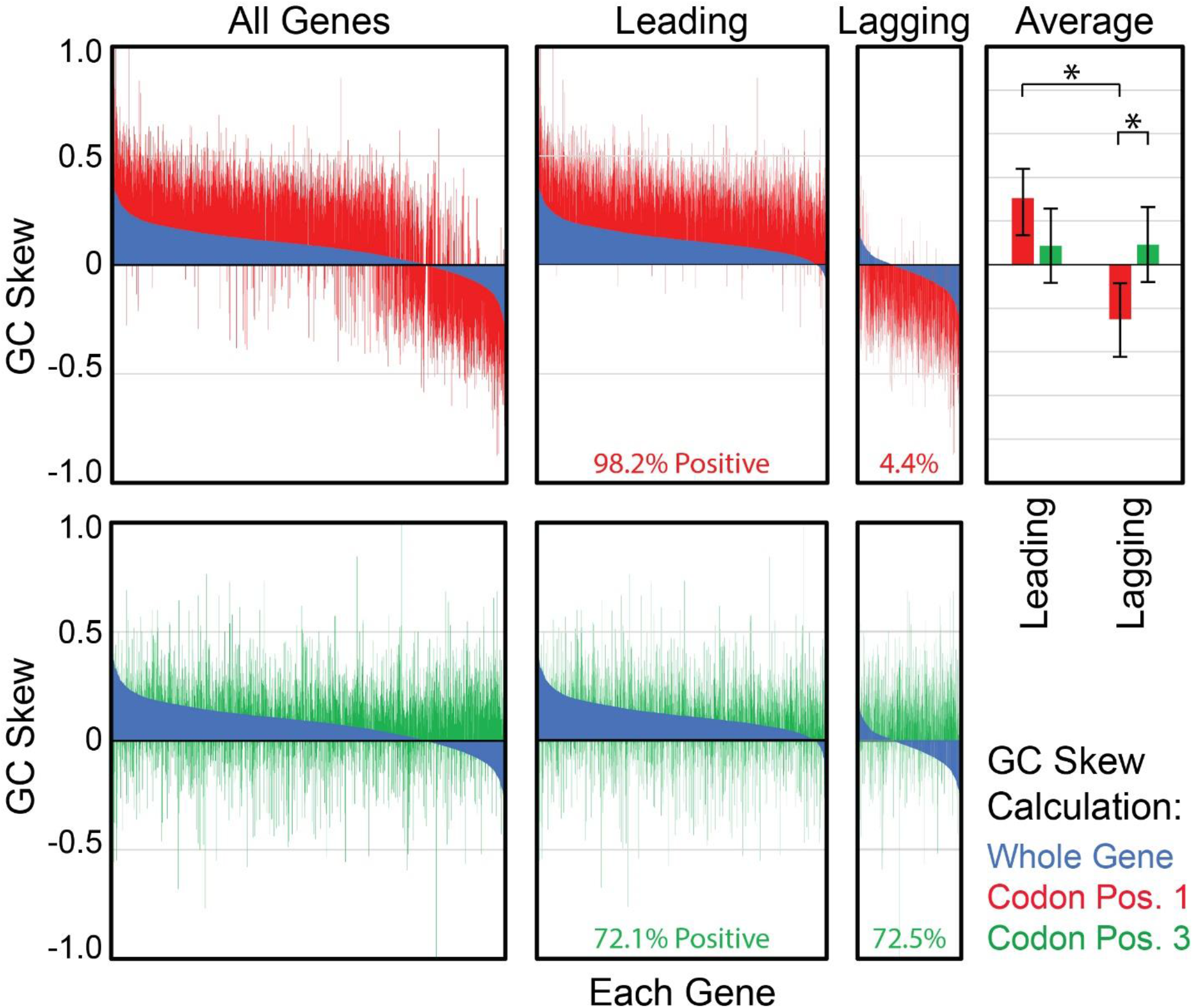
GC Skew values for whole gene regions. Upper graphs: GC skew values are calculated using whole gene regions (blue) or only codon position 1 nucleotides (red) for all genes, leading strand genes only (Leading), or lagging strand genes only (Lagging). Genes are sorted based upon the whole gene GC skew resulting in the appearance of a curve. CP1-based values are not independently sorted, allowing for direct comparison between data sets. The average GC skew values (Average) demonstrate that codon-position 1-based GC skew values (red) are significantly different for leading versus lagging strand genes, whereas codon position 3-based values (green) are not. Additionally, CP1 is significantly different than CP3-based GC skews for lagging strand genes. Error bars represent the standard error of the mean. Significance was determined by the z-test, * indicates p = 0.0. The percentage of positive CP1-based GC skew values is shown at the bottom of the Leading and Lagging strand gene graphs (red text). Lower graphs: Whole gene (Blue, same data as upper graphs) versus codon position 3-based GC skew values are shown. The percentage of positive GC skew values using CP3 nucleotides is identified at the bottom (green text).

To further test the validity of our method and conclusions, we returned to the hypothesis that the CP3-based GC skew value of a GC skew inversion should rise quickly relative to CP1-based values due to lower selective pressure. If this model is correct, then CP3-based GC skew values should generally be higher than CP1-based value. Indeed, that is exactly what we observed (Fig. 2, green versus red columns). This case is particularly clear among lagging strand genes where the average CP3-based value is fully positive while the average CP1-based value remains negative. These differences are highly significant (Z-test p = 0). Notably, these data imply that a CP1-based GC skew analysis should be more accurate than a whole-gene based analysis for detecting genes that change orientation. Therefore, we compared the CP1-based GC skew values to our original data for *B. subtilis* (Fig. 2, red versus blue columns). Incredibly, these data indicate that nearly all (94%) all of the genes currently on the lagging strand were encoded on the leading strand throughout the majority of *B. subtilis*’ evolutionary history. This exceeds our original estimate of 64%. As such, our original data appears to be generally accurate, if limited in resolution (discussed below).

Though most of the differences between the GC skew and phylogeny data sets are due to GC skew reversions, some discrepancies remain. We hypothesized that if the ancestor/descendant pairs diverged long ago, subsequent mutagenesis could have erased the negative value of GC skew inversions. This could explain the remaining data points. In keeping with this notion, previous work shows that the closest related ancestor/descendant pair, *Mycoplasma penetrans* and *Mycoplasma gallisepticum*, diverged at least 100M years ago^3^. (Divergence time is based on 1.8% difference in the 16S rRNA genes between the ancestor and descendant species, and a 1% rate of change per 50M years^3^.) In further support of this temporal limitation hypothesis, we observed that the higher resolution CP1-based GC skew analysis solved the problem: For the *B. lichenformis*/*B. subtilis* pair, a CP1-based GC skew sign change was apparent in 100% of the ortholog pairs (Supplementary Table 1, right columns). To ensure that this does not reflect a high false positive rate, we calculated the CP1 GC skew for all single copy orthologs that did not change orientation (Table S4). In this independent analysis, the two methods show 98.4% agreement. As the remaining discrepancies could reflect an error in either method^4^, our data imply a maximal false negative rate of 1%, and maximal false positive rate of 1.6%. This overwhelming agreement strongly supports the validity of the CP1-based GC skew-based analysis. It also suggests that our original method has a lower temporal limitation that reduces detection of ancient GC skew inversions.

The authors also challenge our inference that positive selection acts more frequently on lagging strand genes. To further test our interpretation, we conducted likelihood ratio tests using the single-nonsynonymous rate models M1a (neutral evolution) and M2a (positive selection) (Figure 2A)^5^. We found that lagging strand genes with a dN/dS value >1 are likely to be under positive selection (Chi square test p<0.05) approximately 2.5 times more frequently than leading strand genes (Fisher’s exact test p = 0.014) in *M. tuberculosis*. We observe a similar ratio in *B. subtilis*, though the difference between the leading and lagging strand genes does not reach 95% significance (Fig. 3, middle). We also repeated our previous cross-species analysis: genes with a dN/dS ratio greater than 1 (Chi square p<0.05) that better fit the M2a model are more frequently observed in lagging strand genes (Fisher’s exact test p = 0.005, Figure 3, right). At the 95% confidence level, we could expect to observe roughly 5 false positive data points among the 108 leading and 97 lagging strand genes with a dN/dS >1. As we observe 19 and 26, respectively, our results are well above background. Therefore, these data suggest that our inference of more frequent positive selection on lagging strand genes was accurate.

**Figure 3.**
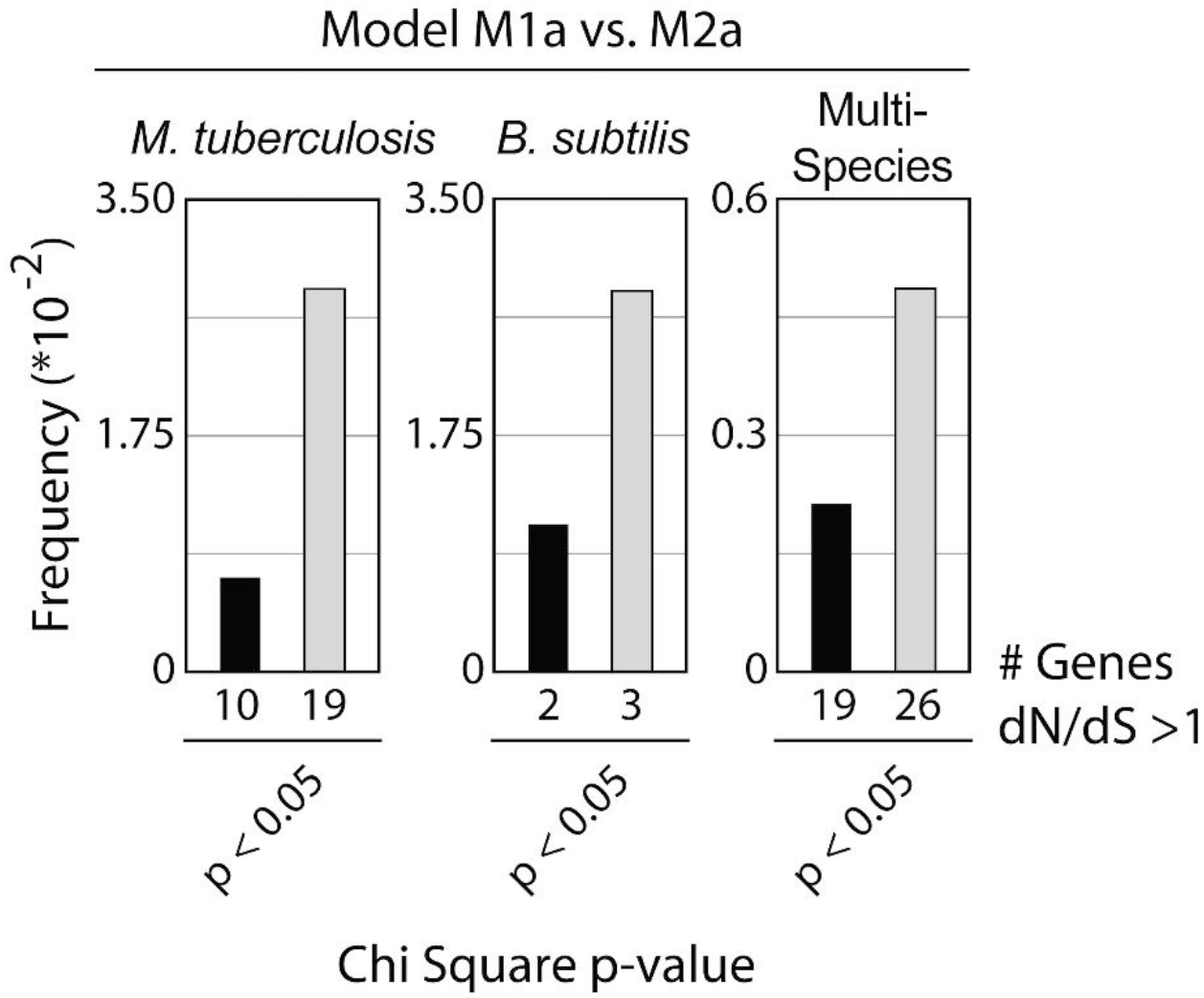
Lagging strand genes are more frequently under positive selection. Likelihood ratio tests (LRTs) were used to compare site models M1a (neutral model) versus M2a (positive selection model) as a test for positive selection among genes with a dN/dS ratio greater than 1. The frequency of genes better fitting the positive selection model at >95% significance are plotted (# genes under positive selection / total). Positive selection likely acts more frequently on lagging strand genes in *M. tuberculosis* (Fisher’s exact test p = 0.049). An equivalent analysis is shown for *B. subtilis* (middle graph, Fisher’s exact test p = 0.12 for leading versus lagging strand frequencies). Multi-species analysis: across species, genes under positive selection are more frequent in lagging strand genes, confirming the results of our original analysis (right graph, Fisher’s exact test p = 0.005).

The authors also claim that the equivalent dS of leading and lagging strand genes indicate that lagging strand genes do not mutate at a faster rate (lines 112-118), and that this metric contradicts the adaptive hypothesis (line 117). This is an interesting point. First, though the dS values in our manuscript are indeed equal for leading and lagging strand genes, the manuscripts cited by the authors directly contradict their claims: both Schroeder et al. and Sankar et al. reported a higher base substitution rate in lagging strand genes (their Figures 3A and 1D, respectively). Likewise, a recent study confirmed the higher mutation rate of lagging strand genes in *E. coli*^6^. Secondly, there are other interpretations of the dS values that do not contradict our model. The dS represents a long-term average mutation rate but is uninformative about short-term variations in spontaneous mutation rates. Our work shows that short-term variations may be a critical concern. We showed that, *when transcribed* at relatively higher levels than the baseline, lagging strand alleles have a higher mutation rate than an otherwise identical leading strand alleles^7,8^. If leading and lagging strand genes were equivalent (expression pattern, gene length, etc.), the authors’ interpretation would be reasonable. However, there are major differences between these groups^1,9,10^. In Figure 4 (top), we show how a leading strand gene and a (distinct) lagging strand gene could accumulate a similar number of mutations over the same time period, while having different mutation rates when transcribed^8,11^. These profiles are based on mutation rates measured using the established using Luria-Delbrück fluctuation assays^8,11^. Here, the lagging strand allele has twice the mutation rate of the leading strand gene when transcribed, but an equal mutation rate when transcriptionally repressed (Figure 4, top). This model predicts that transcriptional induction is a major factor that determines the dS of the lagging strand gene. We also demonstrate how a flipping event will alter the same gene’s mutation rate under an identical induction profile (Figure 4, bottom). Together, these models reconcile the equal dS observed in nature with the higher spontaneous mutation rate of lagging strand alleles observed in highly-controlled laboratory experiments.

**Figure 4.**
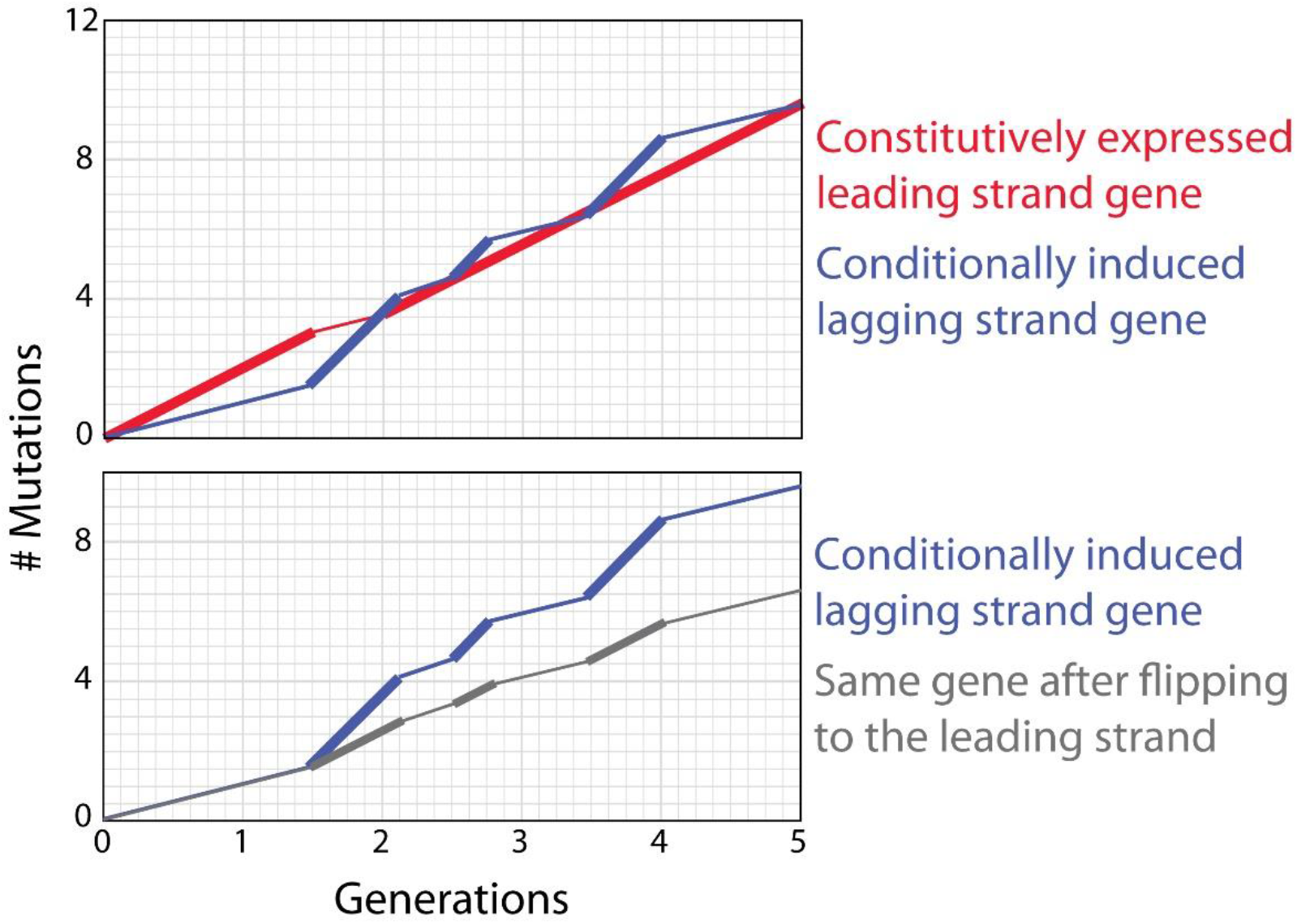
Lagging strand genes can have both a higher mutation rate when transcribed, and also a dS value equivalent to leading strand genes. Upper graph: Comparison of mutation rates for a hypothetical leading strand gene versus a hypothetical (but distinct) lagging strand gene. The two genes depicted have distinct transcriptional induction patterns consistent with observed trends: the leading strand gene is almost always expressed, whereas the lagging strand gene is briefly induced (for 3 short time periods in this example). On a conceptual level, these differences are consistent with patterns commonly observed for genes in each group^16,17^. The number of spontaneous mutations (a proxy for the dS) for each gene is graphed (leading strand gene in red, lagging strand gene in blue). For each gene, thick lines indicate transcriptional induction, and thin lines represent times of transcriptional repression. Mutation rates (the slope of the lines) change depending upon transcriptional activation/repression as previously described, both generally, and for multiple leading vs. lagging strand reporter genes^8,11,18–20^. Here the leading strand gene (Red) gains mutations at a rate of 2 mutations/generation when transcribed, whereas the lagging strand gene (Blue) gains mutations at 4 mutations/generation when transcribed, as observed for the *hisC952* gene^8^. Both genes have a lower and equivalent mutation rate when transcriptionally repressed (1 mutation/generation). These data show that a similar *average* mutation rate (i.e. dS) can be produced by the two genes, despite the higher mutation rate of the lagging strand gene during transcriptional induction. Lower graph: Mutagenesis patterns for the *same* gene when encoded on in the leading versus the laggings strand. This comparison isolates the effects of gene orientation, showing that lagging strand encoding yields a higher mutation rate for the same gene consistent with at least 3 reporter genes^8^. Together the upper and lower graphs can explain how lagging strand encoding increases the mutation rate of individual genes, even while ensemble analyses display equivalent average mutation rates (dS values) for leading and lagging strand genes as a group.

Liu and Zhang further propose that a gene’s GC skew value can become negative due to causes other than gene flipping (GC skew inversion). If correct, this would indeed undermine the validity of our method and conclusions. In support of this notion, the authors reference Chen et al. who showed that transcription, translation, and replication-based nucleotide synthesis cost biases can affect GC and AT skews^12^. Their data suggests that cost bias can increase a gene’s GC skew with respect to the sense strand, resulting in negative GC skew values in lagging strand genes^12^. We hypothesized that the overall mutational bias, including the influence of cost bias, can be identified through our CP1 versus CP3-based GC skew analyses. If the Chen et al. model is correct, the CP3-based GC skew value of lagging strand genes should frequently be lower than the CP1-based values. Alternatively, if our model is correct, DNA replication should universally drive the CP3-based values *higher* than the CP1-based values in lagging strand genes. Our analysis in Figure 2 (green versus red columns) clearly demonstrates that the average CP3-based value is significantly higher than the CP1-based value in *B. subtilis* (Z-test, p = 0). CP3-based values also tend to be positive in absolute terms (Figure 2, Averages). Both observations support our model. However, low GC organisms (e.g. *B. subtilis*) show increased “indifference” toward amino acid related selection^12^. Therefore, we performed the same test in *M. tuberculosis* (strain H37Rv) which has a high GC content. Again, we observed the same pattern: Among the total 1579 lagging strand coding genes, 97% (1531 genes) of CP1-based GC skew values are negative or equal to zero, versus 2% for leading strand genes. We also observe that 81% of lagging strand genes have a positive CP3-based GC skew value. Accordingly, CP3-based GC skew values are higher than the CP1-based value for 95.6% (1510/1579 genes) of lagging strand genes. Even in *E. coli* which has a low strand bias^1^, 89% (1759/1978 genes) of lagging strand genes have a negative CP1-based GC skew value. Among them, the CP3-based GC skew is higher than the CP1-based GC skew in 83% (1463/1759 genes), and 48% (848/1759 genes) of the values are positive. Though this does not directly contradict findings of Chen et al., the data strongly suggest that the net effect of all mutational pressures causes the negative GC skew values of lagging strand genes to increase, equilibrating at a positive value. By extension, gene flipping events appear to be the primary cause of negative GC skew values^1^.

In summary, Liu and Zhang’s technical criticisms are due mainly a misunderstanding over our atypical use of the term “inversion”. After we addressed this semantic conflict and introduced the concept of “GC skew reversions”, it became clear that the authors’ inversion (i.e. “flipping”) data largely agrees with our original (“GC skew inversion”) analysis. Most of the false negative errors the authors identified in our analysis are simply GC skew reversions. However, a small subset of ortholog flips appear to have occurred so long ago (potentially over 100M years ago) that subsequent mutagenesis prevented their detection via the whole-gene GC skew method. These can be considered false negative data points. Alternatively, the positive GC skew of such genes implies that they have acclimatized to the new orientation. Under this perspective, such data points are not false negatives. Either way, we were able to largely resolve this issue by improving the GC skew method through an analysis of first codon position nucleotides. This CP1-based analysis accurately identified an opposing GC skew sign (+/−) in 100% of the authors’ ortholog flips in their *B. licheniformis*/*B. subtilis* comparison, confirming its accuracy. This higher-resolution method allowed us to determine that our original analysis was largely accurate. However, the patterns we identified appear to be far stronger than we initially appreciated: Incredibly, nearly all lagging strand genes may have originally been encoded on the leading strand.

Regarding our inference of increased positive selection on lagging strand genes, we have provided a new statistical analysis of our original dN/dS data. These results also support our original conclusion and the adaptive hypothesis. This is consistent with the observation that convergent mutations (a second indicator of positive selection) also appear to be more common among lagging strand genes^11,13^. Importantly, the authors’ mutation-selection hypothesis cannot explain the convergent evolution data or dN/dS ratios significantly greater than 1. As such, it appears that lagging strand encoding is adaptive for a subset of current lagging strand genes. However, these models are not mutually exclusive. Accordingly, we propose that a combination of neutral evolution and negative selection against highly transcribed lagging strand alleles (the mutation-selection hypothesis), as well as positive selection on a subset of lagging strand alleles, drive the organization of the lagging strand.

## Methods

### Inference of positive selection

We first compiled the nucleotide sequence alignment files produced by TimeZone v.1 and published in our 2018 Nature Communications manuscript. Alignments for genes determined to have a dN/dS value greater than 1 were input into PAML’s codeml application. Likeliood ratio tests of models M1a (null hypothesis) versus M2a (positive selection) were used to determine if each gene is likely under positive selection at the 95% significance level. CodeML settings were based on established standards^14^. The fraction of all leading or all lagging strand genes inferred to be under positive selection were calculated. Fisher’s exact test was used to determine if the observed frequencies are significantly different for leading versus lagging strand genes.

### GC Skew Calculation Method Comparison

The GC Skew, defined as (G−C)/(G+C), was calculated for the leading strand sequence of whole gene regions or using only nucleotides corresponding to the 1^st^, 2^nd^, or 3^rd^ codon positions. The Pearson Correlation coefficient was calculated for the whole gene average method versus the indicated codon position-specific method (Table 1).

### Independent Phylogeny-Based vs. GC skew Analysis

Data are presented in Table S4. The program OrthoFinder was used to identify all single copy orthologs between *B. subtilis* strain 168 and *B. licheniformis* strain ATCC 14580 using default settings. Gene orientations were annotated in Microsoft Excel based upon the replication origins and termini listed on the DoriC database^15^. Codon position 1-based GC skew values were calculated using custom Python scripts.

